# Simultaneous LC-MS determination of glucose regulatory peptides secreted by stem cell-derived islet organoids

**DOI:** 10.1101/2023.06.12.544566

**Authors:** Christine Olsen, Chencheng Wang, Aleksandra Aizenshtadt, Shadab Abadpour, Elsa Lundanes, Frøydis Sved Skottvoll, Alexey Golovin, Mathias Busek, Stefan Krauss, Hanne Scholz, Steven Ray Wilson

## Abstract

For studying stem cell-derived islet organoids (SC-islets) in an organ-on-chip platform, we have developed a reversed phase liquid chromatography tandem mass spectrometry (RPLC-MS/MS) method allowing for simultaneous determination of insulin, somatostatin-14, and glucagon, with improved matrix robustness compared to earlier methodology. Combining phenyl/hexyl-C18 separations using 2.1 mm inner diameter LC columns and triple quadrupole mass spectrometry, identification and quantification were secured with negligible variance in retention time and quantifier/qualifier ratios, negligible levels of carry-over (< 2%), and sufficient precision (± 10% RSD) and accuracy (± 15% relative error) with and without use of internal standard. The here developed RPLC-MS/MS method showed that the SC-islets have an insulin response dependent on glucose concentration, and the SC-islets produce and release somatostatin-14 and glucagon. The RPLC-MS/MS method for these peptide hormones was compatible with an unfiltered off-line sample collection from SC-islets cultivated on a pump-less, recirculating organ-on-chip (rOoC) platform. The SC-islets background secretion of insulin was not significantly different on the rOoC device compared to a standard cell culture well-plate. Taken together, RPLC-MS/MS is well suited for multi-hormone measurements of SC-islets on an organ-on-chip platform.

## 1 Introduction

The development of organoids, i.e. laboratory-grown 3D organ models, is a rapidly growing field with broad implications on biomedical research [1, 2]. By combining organoids with microfluidics in an organ-on-chip (OoC) device, it has been possible to improve aspects of organ functionality *in vitro* compared to static 3D culture systems [3, 4]. Our research group focuses on applying separation science and mass spectrometry technology for studying metabolism of organoids under static conditions and on OoC devices [5, 6].

Pancreatic islets are composed of several endocrine cells, the majority of the cells being insulin-producing beta cells, somatostatin-14-producing delta cells, and glucagon-producing alpha cells [7]. The precise regulation of glucose homeostasis is controlled by the hormone secretion from these cells [8]. In type 1 and type 2 diabetes mellitus, both insulin secretion and glucagon secretion are impaired [9, 10]. Somatostatin-14 is a paracrine inhibitor of the secretion of insulin and glucagon, however, the role of somatostatin-14 in diabetes is not yet fully understood [11].

Stem cell-derived pancreatic islet organoids (SC-islets) are an emerging alternative for disease modeling of diabetes and cell replacement therapy [12, 13]. Similar to natural islets, SC-islets differentiated from embryonic stem cells consist of multiple types of endocrine cells [14, 15]. SC-islets should therefore be suitable for disease modeling and may serve as a source for islet transplantation in beta cell replacement therapy for type 1 diabetes patients [13, 15]. Another important hormone produced in the pancreatic islets is urocortin-3, which is a general maturation marker for both alpha and beta cells in humans [16, 17]. Urocortin-3 would also be beneficial to study in disease modelling, as it is a marker for dedifferentiated beta cells under diabetic conditions (i.e. altered phenotype with loss of insulin capabilities) [17].

The characterization of SC-islets, both under static conditions and on-chip, would benefit from a sensitive and robust methodology for simultaneous determination of multiple hormones. We have previously developed a sensitive reversed phase liquid chromatography – tandem mass spectrometry (RPLC-MS/MS) method for measuring insulin secreted from SC-islets [18]. However, we experienced limitations regarding sample matrix compatibility, i.e. the method was compatible with Krebs buffer (a balanced salt solution used to mimic physiological conditions), but not compatible with cell medium commonly applied in cultivation of organoids. We hypothesized that, by improving the LC separation, the method could be improved to also include other hormones and be suitable for several biologically relevant sample matrices.

We here describe an expanded and more versatile RPLC-MS/MS method for simultaneous determination of insulin, somatostatin-14, and glucagon secreted by SC-islets, and show that the method is compatible with supernatant collected from SC-islets cultivated in a pump-less, recirculating organ-on-chip (rOoC) device [4]. We show that the SC-islets display an insulin response that is dependent on glucose concentrations. In addition, we were able to detect and quantify the release of somatostatin-14 and glucagon from the SC-islets, confirming that the SC-islets contain functional beta-, delta-, and alpha cells.

## 2 Materials and methods

### 2.1 Chemicals and solutions

Acetonitrile (ACN, LC-MS grade), bovine serum albumin (BSA, ≥ 98%), dimethyl sulfoxide (DMSO, ≥ 99.7%), formic acid (FA, 98%), insulin from bovine pancreas (HPLC grade), synthetic glucagon (≥ 95%, human HPLC grade), recombinant insulin human (≥ 98%), somatostatin-14 (≥ 97%, human HPLC grade), and urocortin-3 (≥ 97%, human HPLC grade) were all purchased from Sigma-Aldrich. Taxonomy will only be indicated for insulin, as all other peptides have human origin. Water (LC-MS grade), and methanol (MeOH, LC-MS grade) were obtained from VWR Chemicals (Radnor, PA, USA). Gibco™ basal cell medium MCDB131, GlutaMAX™ supplement (cat. no. 35050061) and minimum essential medium non-essential amino acids (MEM NEAA) stock solution by Gibco™ was acquired from Thermo Fisher Scientific (Waltham, MA, USA).

Krebs buffer was prepared in-house and consists of the following chemicals of analytical grade: 10mM HEPES, 128mM NaCl, 5mM KCl, 2.7mM CaCl_2_, 1.2mM MgSO_4_, 1mM Na_2_HPO_4_, 1.2mM KH_2_PO_4_, 5mM NaHCO_3_, and 0.1% BSA.

Islet maturation cell medium was prepared in-house by adding 1% Penicillin/Streptavidin, 2% BSA, 10 μg/mL of heparin, 10 μM of ZnSO4, 0.1% of trace elements A and B stock solution, 1% of GlutaMAX™ stock solution and 1% of MEM NEAA stock solution to basal MCDB131 cell medium.

### 2.2 Preparation of individual hormone solutions and calibration solutions

Aqueous water solutions of human insulin, somatostatin-14, glucagon, urocortin-3, and bovine insulin (applied as internal standard for human insulin) were prepared individually by dissolving 1 mg of peptide powder in 1 mL of 0.1% FA in water. The 1 mg/mL stock solutions were further diluted to working solutions consisting of 10 ng/µL of each individual peptide, and divided into 100 µL aliquots. Aliquots of human and bovine insulin were kept at −20 °C until use or for a maximum of three months. Aliquots of somatostatin-14, glucagon, and urocortin-3 were kept at −80 °C until use. All solutions containing proteins were prepared in protein low binding tubes from Sarstedt (Nümbrecht, Germany).

Separate standard solutions of 10 ng/µL of somatostatin-14, glucagon, and urocortin-3 in a 1+1 mixture of ACN and water were prepared for direct injections on the MS. Solutions with human and bovine insulin in Krebs buffer and islet maturation cell medium were prepared in the same manner as described for water-based solutions, with the exception being the amount of FA: 0.5% FA in Krebs buffer and 1% FA in cell medium. Krebs buffer and cell medium solutions were spiked with separate water-based solutions of 10 ng/µL somatostatin-14, glucagon, and urocortin-3 and further diluted with the appropriate matrix to obtain the desired concentrations.

For assessment of the LC-MS method and the preparation of calibration solutions, freshly thawed working solutions of the peptides were further diluted to the desired pg/µL concentration with the experiment appropriate matrix, and spiked with bovine insulin to a concentration of 5 pg/µL unless stated otherwise. Quality controls (QC) were prepared in the same manner.

### 2.3 Cell culture, differentiation, and glucose stimulated insulin secretion for stem cell-derived islets

The SC-islets examined in this study, are prepared according to a previously described differentiation protocol [18]. In brief, SC-islets were generated from the human pluripotent cell line H1 (WA01, WiCell, Madison, WI, USA) with a stepwise differentiation protocol. Following the differentiation, the cells were aggregated as spheroids on an orbit-shaker for 7 days prior to analysis. For static conditions in Krebs buffer, batches of 30 SC-islets were placed in 24-well cell culture plates, hormone secretion was assessed by exposure to: 1 mL Krebs buffer with 2 mM glucose for 60 min at 37 °C, 1 mL Krebs buffer with 20 mM glucose for 60 min at 37°C, and 1 mL Krebs buffer containing 20 mM glucose plus 30 mM KCl for 30 min at 37 °C. Up to 900 µL of supernatant was collected.

Insulin in the supernatant was quantified with human insulin enzyme-linked immunosorbent assay (ELISA) kit (Mercodia, Uppsala, Sweden) and with the LC-MS/MS method described in this study. Prior to injection on the LC-MS/MS system, the collected supernatants were spiked with a total of 5 pg/µL of bovine insulin and 100% FA was added to a total of 0.5%.

### 2.4 Adsorption experiments on a poly(methylmethacrylate)-based organ-on-chip platform

Experiments were performed on the poly(methylmethacrylate) (PMMA)-based rOoC platform, which was fabricated in-house using laser-cutting and thermobonding as previously described in [4]. The rOoC platform consists of two nested circuits of perfusion channels separated with two organoids chambers. Channels are separated from the organoid chambers by a step reservoir with a height of 300 µm.

To evaluate surface adsorption in the rOoC platform, a solution consisting of 2 ng/µL of human insulin in Krebs buffer was incubated on-chip and in standard cell culture 24-wells plate from Corning (Corning, NY, USA) for 20 hours, and compared to an aliquot stored in the freezer.

### 2.5 Background secretion experiments from stem cell-derived islets cultivated on-chip and in static system

SC-islets were cultured in islet maturation cell medium in a rOoC platform with a modified design compared to [4], where the chamber for SC-islets was placed under the perfusion channel to facilitate media and oxygen exchange. In the organoids chamber, batches of 3-6 SC-islets were embedded in extracellular matrix (Geltrex, Gibco, cat. n A1569601) and cultured for 7 days under perfusion. The cell medium in the channels was collected after 24 h of incubation on day 5 and day 7. In the off-chip control culture, batches of 14-19 SC-islets were cultured in 24-well plates for 7 days, and cell medium was collected after 24 h of incubation on day 5 and day 7.

### 2.6 Gel electrophoresis

A water-based solution consisting of 10 ng/µL human insulin and 10 ng/µL BSA was used as a protein marker during gel electrophoresis. To 30 µL of the sample, 10 µL of Bolt™ LDS sample buffer (4x) was added prior to heating at 70 °C for 10 min on a Thermo-Shaker from Grant instruments (Shepreth, UK). The samples were loaded with ultrafine pipettes, from VWR, onto a Bolt™ 4-12% 2-(bis(2-hydroxyethyl)amino)-2-(hydroxymethyl)propane-1,3-diol (bis-tris) plus gel inserted in a mini gel tank. The chamber was filled with Bolt™ 2-(N-morpholino)ethanesulfonic acid (MES) SDS running buffer (20x), diluted to 1x with water. All Bolt™ products and the mini gel tank were from Thermo Fischer Scientific. For 20 min, a voltage of 200 V was applied to the gel by a power supply from Delta Elektronika (Zierikzee, Netherlands). Subsequently, the gel was washed four times with water for five minutes on a shaker. Following the wash, the gel was covered with Imperial™ protein stain (from Thermo Fischer Scientific) and left on shaking for 15 min. Before photographing the gel using a smartphone camera, the gel was washed with water four times for five minutes, followed by washing with gentle shaking in water overnight (18 hours).

### 2.7 Liquid chromatography - mass spectrometry instrumentation

The LC-MS system applied in this study has been used for insulin determination previously [18]. In brief, the conventional LC system was a modified Agilent 1100 series pump (Santa Clara, CA, USA) employing only shielded fused silica connectors (shielded fused silica nanoViper™ sheated in polyetheretherketone tubing from Thermo Fisher). Injection was achieved by coupling a 6-port-2-position valve, with a 50 µm inner diameter (id) x 550 mm (1.08 µL) shielded fused silica loop or a 20 µL shielded fused silica loop. A 250 µL glass syringe was used for injection, following injection the syringe and the loop was washed by flushing the syringe three times with 50/50 MeOH/water, before washing the loop twice with the MeOH/water solution. Prior to next injection, the syringe was washed once with 0.1% FA in water and the loop was washed twice with 0.1% FA in water. The column set-up consisted of an Accucore™ phenyl/hexyl guard column (2.1 mm id x 10 mm, 2.6 µm, 80 Å) attached within a Uniquard drop-in holder to the InfinityLab Poroshell EC-C18 separation column (2.1 mm id x 50 mm, 2.7 µm, 120 Å).

The applied MS was a TSQ Quantiva triple quadrupole MS equipped with a heated electrospray ionization source (H-ESI-II probe) both from Thermo Fisher. The vaporizer temperature was set to 210 °C, and a spray voltage of 3.5 kV was applied to the H-ESI-II probe. Sheath gas was set at 20 Arb (approx. 2.7 L/min), while auxiliary gas was set at 9 Arb (approx. 7.5 L/min), and sweep gas was not applied. The ion transfer tube temperature was kept at 275 °C.

### 2.8 Optimized gradient settings for LC separation

The mobile phase reservoirs contained 0.1% FA in water added 1% DMSO and 0.1% FA in ACN added 1% DMSO, respectively. A 150 µL/min solvent gradient was started at 1% B, quickly increased to 25% B in 1 min, then linearly increased to 32.5% B in 6 min, and kept at 32.5% B for 4 min. In the following step, the %B was quickly increased to 80% B and kept at 80% for 2 min, before quickly decreased to 40% B, and kept at 40% B for 3 min, before being further decreased to 1% B and kept at 1% B for 7 min. The gradient had a total runtime of 23 min, including column re-equilibration for 7 min at 1% B.

### 2.9 Alternative guard and separation column

Other columns examined in this study included an Accucore™ C18 guard column (2.1 mm id x 10 mm, 2.6 µm, 80 Å) from Thermo Fisher, and a Cortecs Premier C18 separation column (2.1 mm id x 10 mm, 2.7 µm, 90 Å) from Waters (Milford, MA, USA).

### 2.10 Settings for selected reaction monitoring

The collision energies and radio frequency (RF) lens voltage were optimized using the compound optimization provided in Xcalibur. The transitions, used in selected reaction monitoring (SRM), including collision energies and RF lens voltage, are listed in **Table 1**. The collision gas (argon) pressure in q2 was 4.0 mTorr and the cycle time was set to 1 sec (equal to 77 ms or 91 ms dwell time per transition with and without urocortin-3, respectively).

**Table 1:**
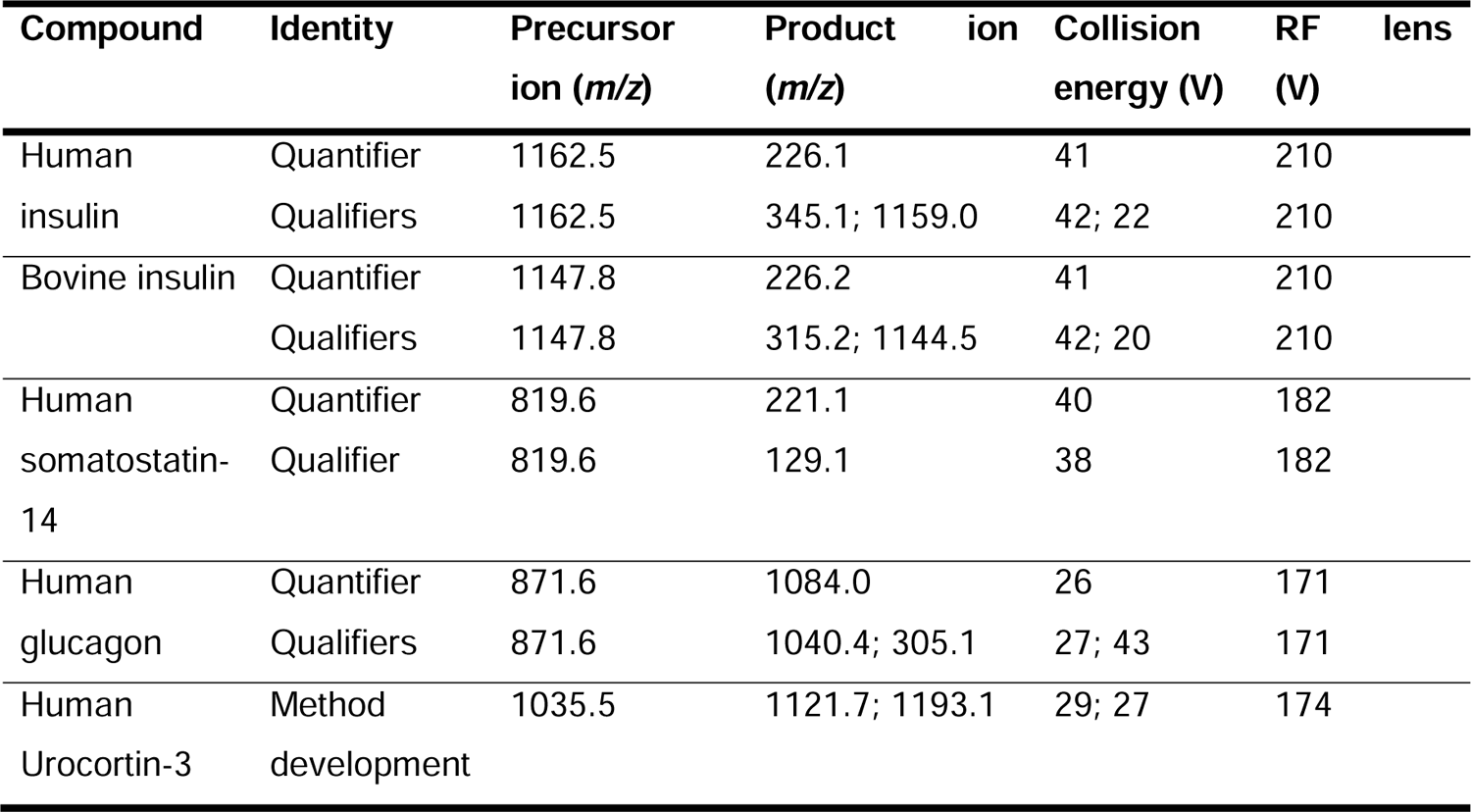
SRM transitions used for human insulin, bovine insulin (internal standard for human insulin), human somatostatin-14, human glucagon, and human urocortin-3 including quantifier/qualifier status, precursor ion, product ion, and collision energy. The settings applied for human urocortin-3 during method development are also included.

## 3 Results and discussion

### 3.1 Optimized gradient separation allowed for simultaneous determination of insulin, somatostatin-14 and glucagon in bovine serum albumin rich matrices

Initial MS/MS settings for somatostatin-14, glucagon, and urocortin-3 were established by direct injection: *m/z* 819.6 ➔ *m/z* 221.1, 129.1 (somatostatin-14), *m/z* 871.6 ➔ *m/z* 1084.0, 1040.4, 305.1 (glucagon), and *m/z* 1035.5 ➔ *m/z* 1121.7, 1193.1 (urocortin-3). The MS/MS settings for human and bovine insulin were selected previously to be: *m/z* 1162.5 (+5) ➔ *m/z* 226.1, 345.0, 1159.0 and *m/z* 1147.8 (+5) ➔ *m/z* 226.2, 315.2, 1144.5, respectively [18]. The four analytes and the internal standard were successfully separated in a water-based solution with the originally applied gradient (8 min separation window from 1% B to 60% B [18]), shown in **Figure 1A**. However, when the analytes were dissolved in Krebs buffer, the separation of urocortin-3 from BSA was not possible, see **Figure 1B**. BSA is the main component in both Krebs buffer and in the cell medium that is used for culturing SC-islets, containing 0.1% BSA and 2% BSA, respectively. BSA produces a multitude of interfering peaks in the *m/z* range from 1000-1500 [18]. Effort was put towards optimization of the gradient, in order to achieve better separation of the peptide hormones from BSA.

**Figure 1:**
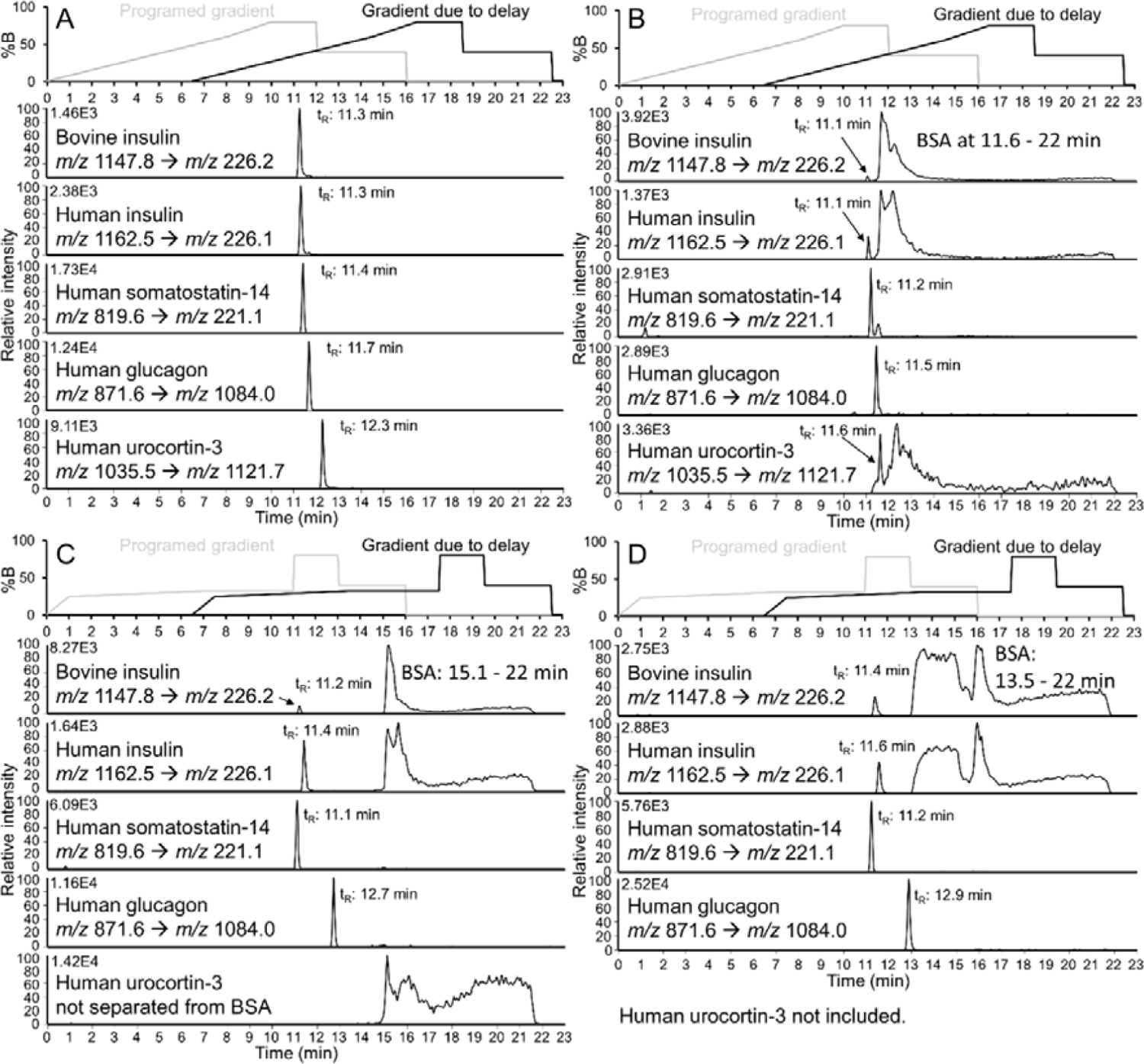
Separation of a peptide mix consisting of bovine insulin, human insulin, somatostatin-14, glucagon and urocortin-3 in different sample matrices. With the original steep gradient: (**A**) 1.08 µL injection of 250 pg/µL peptide mix in 0.1% FA in water, and (**B**) 20 µL injection of 2 pg/µL peptide mix in 0.5% FA in Krebs buffer. With an optimized shallow gradient: (**C**) 20 µL injection of 5 pg/µL peptide mix in 0.5% FA in Krebs buffer, and (**D**) 20 µL injection of 5 pg/µL peptide mix in 1.0% FA in cell medium. The first panel above each separation shows the programed gradient and the gradient due to system delay. The system delay of 6.5 min in the gradient delivery was estimated by running the analysis under non-retained conditions starting at 60% B, see **SI-2** for more details.

By applying a shallow gradient from 25% B to 32.5% B, with an 1.25% B increase per minute, followed by an isocratic step at 32.5% B for 4 minutes, the BSA interference was removed from the insulins, somatostatin-14, and glucagon in both Krebs buffer (**Figure 1C**) and cell medium (**Figure 1D**). Urocortin-3 remained impossible to separate from the BSA in the Krebs buffer in the current set-up, as shown in **Figure 1C**, and was not examined further in this study (see Conclusions sections for further discussion). In addition, the collision gas pressure was increased from 2.5 mTorr used in the original method to 4.0 mTorr due to significantly increased peak area for human insulin and glucagon, see **SI-1** for more details.

In conclusion, by optimizing the applied gradient in the LC separation, the existing RPLC-MS/MS method could be improved to include three of the main hormones produced in islets: Insulin, somatostatin-14, and glucagon, while not being limited by the sample matrix. However, the current column set-up was not suitable for the inclusion of a more retained hormone, urocortin-3, which was not separated from the matrix components.

#### 3.1.1 Guard cartridge: not one-size-fits-all for determination of intact peptides

In the previous section and in Olsen et al. [18], an Accucore phenyl/hexyl guard cartridge (80 Å, 2.6 µm particles, 2.1 mm inner diameter x 10 mm) combined with a Poroshell EC-C18 separation column (120 Å, 2.6 µm particles, 2.1 mm inner diameter x 50 mm) were applied in the LC-MS system. Traditionally, the guard column is packed with the same particles and stationary phase as the separation column. However, it is commonly accepted that combining different stationary phases on a two-column set-up, with a trapping column and a separation column, can improve the separation [19]. The combination of an Accucore phenyl/hexyl trapping column and Poroshell C18 separation column has previously been successfully applied for determination of insulin in urine matrix [20].

To examine in detail if the phenyl/hexyl guard cartridge significantly affected the separation, the separation was compared with and without the phenyl/hexyl guard. In addition, an equivalent Accucore C18 guard cartridge (80 Å, 2.6 µm particles, 2.1 mm inner diameter x 10 mm) was assessed as an alternative to the phenyl/hexyl guard. Asymmetry factors and peak areas obtained on different column set-ups for the peptides are summarized in **Table 2**.

**Table 2:** Chromatographic performance by injection of a peptide mix consisting of 5 pg/µL of bovine and human insulin, somatostatin-14, and glucagon. The obtained asymmetry factor (A_s_) and peak area of the peptides are shown for the following guard cartridge options: Accucore phenyl/hexyl, Accucore C18, and no guard cartridge, combined with the Poroshell EC-C18 separation column (N = 3).

| Columns | Accucore phenyl/hexyl-Poroshell C18 | Accucore C18-Poroshell C18 | No guard-Poroshell C18 |
| --- | --- | --- | --- |
| **Bovine insulin** |  |  |  |
| $A_s$ | 1.9 (14% RSD) | 4.6 (13% RSD) | 2.2 (13% RSD) |
| Peak area | $8.64 \times 10^3$ (13% RSD) | $4.92 \times 10^3$ (6% RSD) | $5.63 \times 10^3$ (6% RSD) |
| **Human insulin** |  |  |  |
| $A_s$ | 2.3 (14% RSD) | 5.6 (7% RSD) | 1.8 (13% RSD) |
| Peak area | $7.79 \times 10^3$ (6% RSD) | $6.71 \times 10^3$ (3% RSD) | $8.05 \times 10^3$ (5% RSD) |
| **Somatostatin-14** |  |  |  |
| $A_s$ | 1.5 (2% RSD) | 1.7 (19% RSD) | 1.4 (4% RSD) |
| Peak area | $1.70 \times 10^4$ (10% RSD) | $1.99 \times 10^4$ (2% RSD) | $2.30 \times 10^4$ (4% RSD) |
| **Glucagon** |  |  |  |
| $A_s$ | 1.5 (22% RSD) | 2.1 (10% RSD) | 1.8 (8% RSD) |
| Peak area | $5.97 \times 10^4$ (4% RSD) | $4.44 \times 10^4$ (3% RSD) | $4.57 \times 10^4$ (1% RSD) |

The asymmetry factor obtained for human insulin increased from 2.3 (N = 3, technical replicates on LC-MS), **Figure 2A**) with the phenyl/hexyl guard to 5.6 (**Figure 2B**) on the C18 guard cartridge, and the peak area was reduced from 7.79 x 10^3^ on the phenyl/hexyl to 6.71 x 10^3^ on the C18. Without applying a guard cartridge, the asymmetry factor for human insulin was 1.8 (**Figure 2C**) and the peak area was 8.05 x 10^3^. By one-factor analysis of variance (ANOVA), there was no significant difference in the obtained peak area of human insulin with or without guard cartridge or type of guard. However, the tailing effect for human insulin obtained on the C18 guard was significantly larger compared to that on the phenyl/hexyl guard or without applying a guard. Additionally, there was no signal detected of human insulin in a subsequent blank injection on the set-up without guard or with the phenyl/hexyl guard, but the C18 guard contributed to 5% carry-over of human insulin.

**Figure 2:**
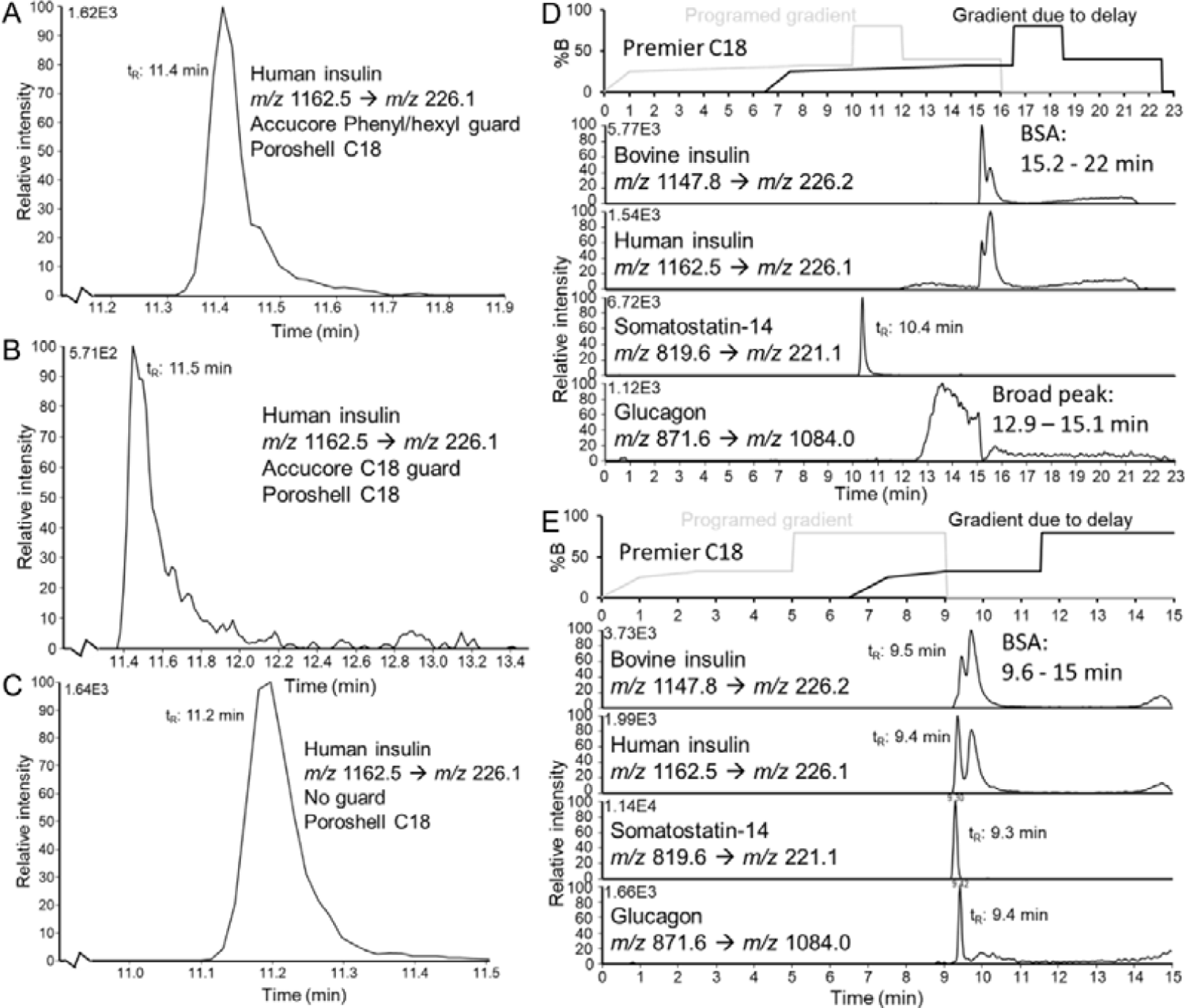
Comparison of peak shape obtained for human insulin on the following column set-ups (**A**) Accucore phenyl/hexyl guard – Poroshell C18, (**B**) Accucore C18 guard – Poroshell C18, and (**C**) only the Poroshell C18 column. Attempted separation of 5 pg/µL peptide mix in 0.5% Krebs buffer on the Premier column with different gradients: (**D**) Insufficient separation of insulins and glucagon from BSA with the same gradient as applied for the Poroshell Column. (**E**) Following attempted gradient optimization, still insufficient separation of insulins and BSA, and low signal intensity for glucagon.

For bovine insulin, the asymmetry factor was not significantly different for the various column set-ups, however the peak area was significantly higher with the phenyl/hexyl guard compared to the C18 guard. The peak area obtained with the phenyl/hexyl guard was not significantly different to the peak areas obtained without guard.

For somatostatin-14, the asymmetry was not significantly different for the different column set-up, however, the peak area was significantly increased without a guard cartridge being applied compared to the phenyl/hexyl guard, information summarized in **Table 2**. There was no significant difference between the C18 and the phenyl/hexyl guard or C18 and without guard applied concerning peak areas obtained for somatostatin-14.

For glucagon, the peak areas were significantly higher when applying the phenyl/hexyl guard cartridge compared to no guard cartridge applied or the C18 cartridge, while the asymmetry factor was not significantly different for the different column set-ups, summarized in **Table 2**.

The differences in the obtained chromatographic performances show that there isn’t a “one-size-fits-all” approach when selecting guard cartridge for determination of various intact peptides, and large effects for asymmetry factor and peak areas can be seen based on changing the stationary phase. Based on our findings, it is hard to conclude why the C18 guard cartridge caused dramatic tailing for human and bovine insulin, and if there was an effect of having particles with smaller pore sizes in the guard cartridge compared to the separation column (80 Å vs 120 Å). When considering the use of guard cartridge, the LC-MS method was intended for experiments with supernatant collected directly from SC-islets possibly containing cell debris and particles, which would not be beneficial to inject directly onto the more expensive separation column without the protection from a guard cartridge.

In conclusion, we continued to employ a phenyl/hexyl guard cartridge based on the significantly higher peak areas obtained for glucagon and the guard cartridge’s protection of the separation column when analyzing unfiltered samples collected directly from SC-islets.

#### 3.1.2 Success of intact hormone analysis is dependent on type of separation column

All three hormones and the internal standard have a minimum of one sulfur-containing amino acid residue (i.e. cysteine and methionine), which may cause adsorption on stainless steel column housing and contribute to band broadening [21, 22]. To see if this was the case in out set-up and if the chromatographic performance could be improved, a Premier column (C18, 2.6 µm, 90 Å) with modified metal surfaces in the column housing and filters to reduce non-defined adsorption was compared to the Poroshell column employed in previous sections [23]. For our particular application, the peaks of insulins and glucagon obtained on the Premier column were broader and not sufficiently separated from BSA or other matrix components in Krebs buffer, **Figure 2D**. The lack of separation indicated that the gradient optimized for the Poroshell column (C18, 2.7 µm, 120 Å) was not suitable for the Premier column. In the attempt to optimize the separation, the peak area of glucagon was greatly reduced for each subsequent injection (from a peak area of approx. 2 x 10^5^ to a peak area < 1 x 10^4^, results not shown), and none of the insulins were successfully separated from BSA, **Figure 2E**. On the Accucore phenyl/hexyl-Poroshell C18 column set-up, an equivalent injection would give a stable peak area of glucagon around 250000. For somatostatin-14, the Premier column set-up was able to provide equivalent chromatographic performance as the phenyl/hexyl-Poroshell C18 column set-up.

Based on the lack of separation from BSA and the loss of signal for glucagon on the Premier column, the Premier column was not found appropriate for our application and was not examined further. One of the major differences between the Premier column and the Poroshell column is the pore size of the particles of 90 Å and 120 Å, respectively. In the previous **Section 3.1.1**, the guard cartridges with 80 Å particle pore size was sufficient for sample loading. However, from the comparison of the Premier and Poroshell column, it is clear that larger pores in the separation column was necessary to be able to separate the intact hormones from BSA and other matrix components [18, 24].

In conclusion: of the columns tested, the originally applied Poroshell C18 separation column was found to be the best option for the simultaneous determination of the three peptide hormones in complex matrices.

### 3.2 Satisfactory accuracy and precision in determining concentration of insulin, somatostatin-14, and glucagon in quality controls in Krebs buffer

Glucose stimulated insulin secretion is a standard characterization experiment of insulin response in SC-islets using Krebs buffer [25, 26]. The experiments require highly accurate determinations of hormone concentrations in Krebs buffer. Therefore, the method’s repeatability concerning linearity, precision, and accuracy in determination of the concentration of the three hormones was examined by establishing a linearity curve in the range from 5 pg/µL to 15 pg/µL in Krebs buffer over three days. The precision and accuracy for determination of hormone concentrations were evaluated based on concentration found in QCs containing 10 pg/µL of each analyte.

The established linearity curves for human insulin (with bovine insulin as internal standard, see **Figure 3A**), somatostatin-14 (**Figure 3C**), and glucagon (**Figure 3E**) did not show sign of heteroscedasticity (i.e. difference in variation of the response depending on the concentration level) as the residuals appear to fall randomly around the x-axis in the residual vs concentration plots in **Figure 3B**, **Figure 3D**, and **Figure 3F**, respectively [27]. The hypothesis of homoscedasticity was confirmed with an F-test comparing the variance in the response of the standard with the lowest and highest concentration levels for each of the analytes.

**Figure 3:**
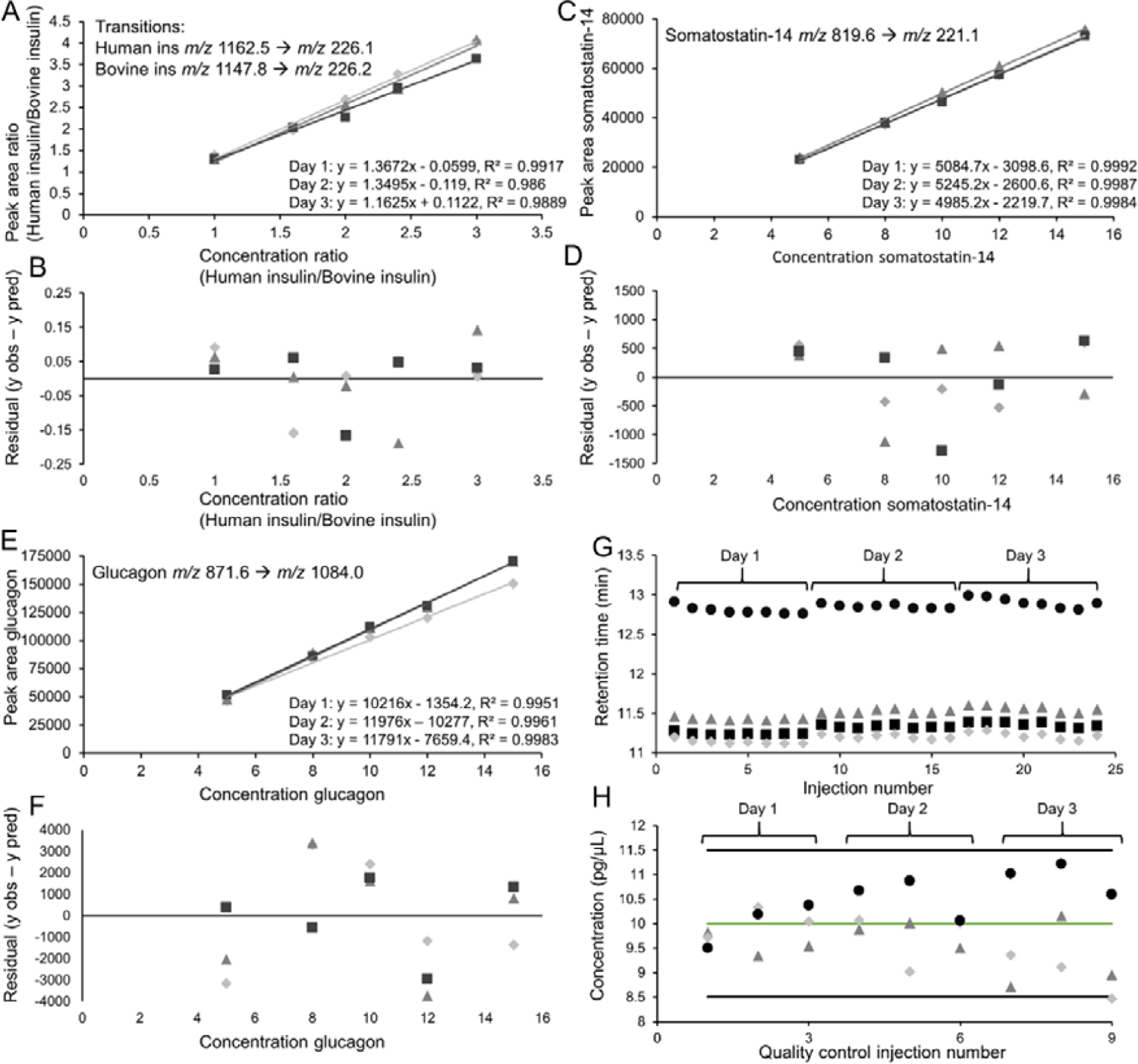
Established linearity curves in the range from 5 pg/µL to 15 pg/µL for (**A**) human insulin with bovine insulin as internal standard, (**C**) somatostatin-14, and (**E**) glucagon, where day 1 is represented with diamonds, triangles for day 2 and squars for day 3. Residual vs concentration plots for each curve belonging to (**B**) human insulin, (**D**) somatostatin-14, and (**F**) glucagon, where day 1 is represented with diamonds, triangles for day 2 and squars for day 3. (**G**) The retention time for bovine insulin (squares), human insulin (triangles), somatostatin-14 (diamonds), and glucagon (circles) over the 24 injections of calibration standards and QCs. (**H**) Determined concentration in QCs for human insulin (triangles), somatostatin-14 (diamonds), and glucagon (circles) in the nine injections.

Bovine insulin was a suitable internal standard for human insulin due to similar structures, similar retention time and the variation of the peak area of bovine insulin was 4% RSD (N = 5) on day 1, 3% RSD (N = 5) on day 2, and 3% RSD (N = 5) on day 3. The variation in the peak area of bovine insulin was much smaller during these examinations compared to the examination done in **Section 3.1.1**, as changes to the system was not done between injections allowing for a better stability. Determination of somatostatin-14 and glucagon was done without use of bovine insulin as internal standard, due to large differences in the structures of the analytes compared to bovine insulin, and different retention time. These differences indicates that bovine insulin is not a suitable internal standard as it cannot compensate for differences in matrix effect (different retention time) nor the transfer from ions in solution into gas phase occurring during the ESI process (different structures). Bovine insulin could compensate for differences in injection volume, however, there is relatively small variation between repeated injections that such compensation was not deemed necessary. The retention time variance over the three days was ≤ 0.5% RSD (N = 24) for all of the hormones, see **Figure 3G**.

In determination of QC analyte concentration using the same standard solutions to establish a calibration curve, the accuracy was within ± 10% relative error (N = 3, per day), ± 10% RSD (N = 3) intra-day precision and ± 10% RSD (N = 9) inter-day precision. Determination based on a single injection of the QC, only somatostatin-14 was not determined with an accuracy within ± 15% relative error, as shown in **Figure 3H** for injection number nine. The average concentration of each analyte found in the QCs on the three separate days was not significantly different determined with one-way ANOVA.

To summarize, the RPLC-MS/MS method, featuring a phenyl/hexyl guard and Poroshell C18 separation column combined with triple quadrupole mass spectrometry, was successful in simultaneous determination of the three analytes; human insulin (including bovine insulin as internal standard), somatostatin-14, and glucagon in Krebs buffer with sufficient accuracy, precision, and repeatability.

### 3.3 Stem cell-derived islet organoids show glucose concentration dependent insulin response and potassium dependent release of somatostatin-14 and glucagon in Krebs buffer

As mentioned in the previous section, the secretion of peptide hormones in SC-islets can be examined by exposing the SC-islets to Krebs buffer containing various amounts of glucose. It has also been shown that the three major cell types found in islets, beta-, delta-, and alpha cells, release a large pool of stored hormones when exposed to high levels of potassium due to direct membrane depolarization [11, 28, 29]. Hence, to show compatibility between supernatant collected from SC-islets in Krebs buffer and RPLC-MS/MS, we attempted to measure the three hormones in Krebs buffer collected from SC-islets exposed to: (1) low amount of glucose (2 mM), (2) high amount of glucose (20 mM) and (3) a combination of 20 mM glucose and 30 mM KCl.

The concentration of insulin determined with RPLC-MS/MS in the supernatant collected from SC-islets challenged with low glucose (**Figure 4A**) was on average 2.2 pg/µL (RSD = 13%, n = 4 batches of SC-islets, N = 1), while there was 4.1 pg/µL insulin (RSD = 10%, n = 4, N = 1) secreted by SC-islets challenged with high glucose (**Figure 4B**). The stimulation index of insulin secretion during the glucose challenge was 1.86 (RSD = 5%, n = 4, N = 1). The amount of insulin determined in the samples collected with 20 mM glucose and 30 mM KCl (**Figure 4C**) was 18 pg/µL (RSD = 14%, n = 4, N = 1), which was eight times higher than the response in low glucose. Simultaneously, the reliability of the determination of insulin using the RPLC-MS/MS method was supervised by QCs, to see if the reported concentrations was within 15% relative error. QCs were included at 2 pg/µL (N = 2), 8 pg/µL (N = 3) and 18 pg/µL (N = 2), and the insulin concentration was determined within 15% relative error for each injection.

**Figure 4:**
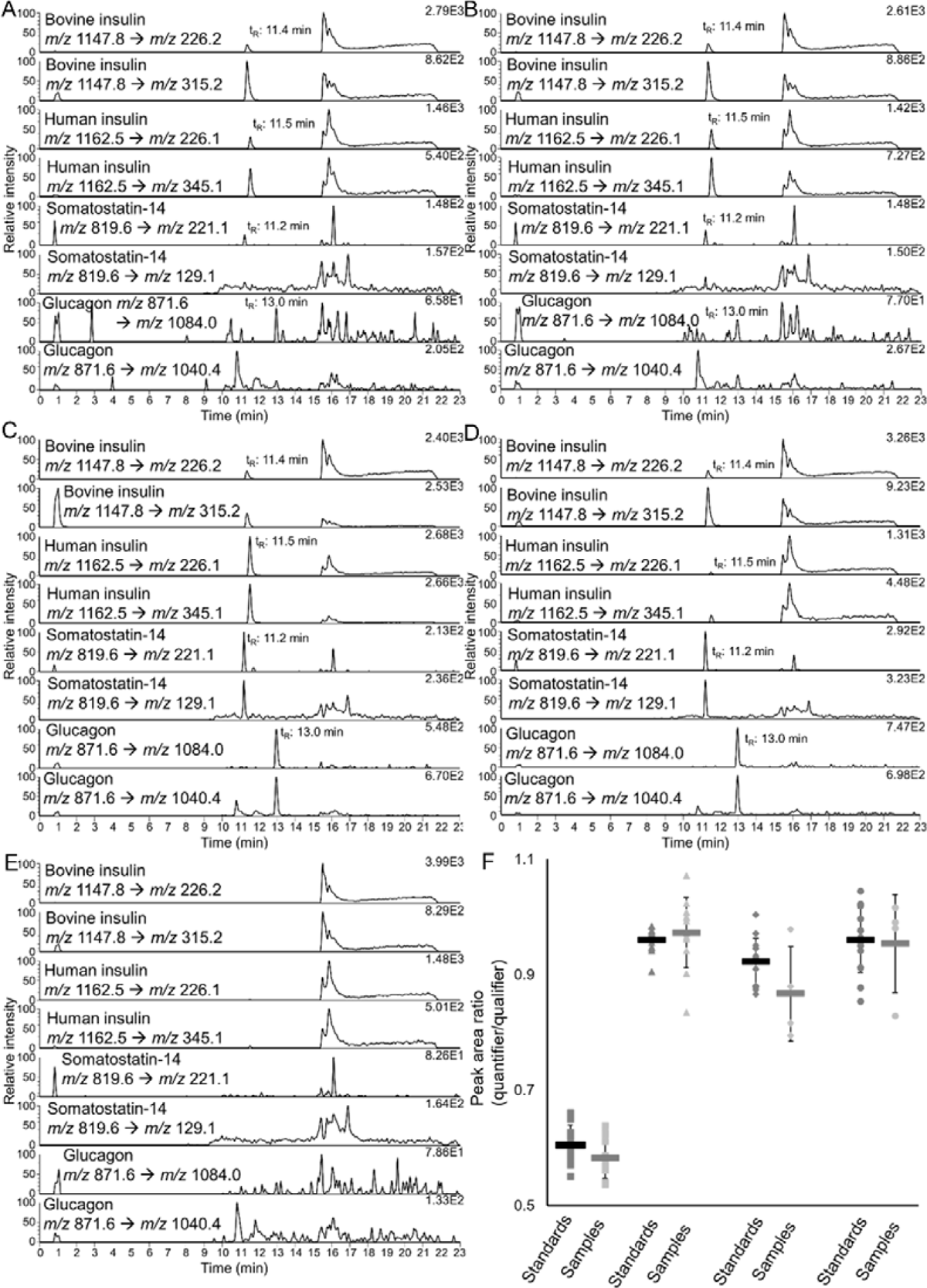
Representative chromatograms obtained in SRM mode of: Supernatant from SC-islets exposed to (**A**) 2 mM glucose, (**B**), 20 mM glucose, and (**C**) 20 mM glucose with 30 mM KCl. (**D**) Calibration standard with 0.25 pg/µL human insulin, somatostatin-14, and glucagon with 5 pg/µL bovine insulin in 0.5% FA in Krebs buffer and (**E**) blank injection of 0.5% FA in Krebs buffer. The first transition for each hormone is the quantifier, while the second transtion is the qualifier. (**F**) The ratio of quantifier/qualifier transitions for bovine insulin (squares), human insulin (triangles), somatostatin-14 (diamonds), and glucagon (circle) obtained for calibration standards and QCs (dark grey), and samples (light grey).

The insulin concentration in the supernatant collected from the SC-islets was also determined with an established ELISA method, which found the following insulin concentrations: 2.6 pg/µL (low glucose, RSD = 5%, n = 4, N = 1), 4.5 pg/µL (high glucose, RSD = 13%, n = 4, N = 1), and 20 pg/µL (KCl, RSD = 13%, n = 4, N = 1). An independent two sample t-test, at 95% confidence, showed that the concentrations determined by LC-MS/MS were not significantly different from the concentrations determined with ELISA.

The same samples were also simultaneously examined for somatostatin-14 and glucagon. Neither the quantifier nor the quantifier transition of somatostatin-14 and glucagon was detected in the supernatant from SC-islets exposed to low (**Figure 4A**) or high glucose (**Figure 4B**), however, in supernatant collected after exposure to KCl, detectable signals were obtained for both hormones and transitions (**Figure 4C**). For somatostatin-14, the average peak area was 1.3 x 10^3^ (RSD = 7%, n = 4, N = 1) after KCl exposure, which was lower than the peak area of 1.9 x 10^3^ obtained for the calibration standard with the least amount of somatostatin-14 of 0.25 pg/µL (**Figure 4D**). For glucagon, the concentration was determined to be 0.28 pg/µL (RSD = 18%, n = 4, N = 1) after KCl exposure. The QCs examined for somatostatin-14 and glucagon at 2 pg/µl and 8 pg/µL were determined within 11% relative error (RSD < 10 %), showing that the RPLC-MS/MS method for somatostatin-14 and glucagon has sufficient precision and accuracy without use of an internal standard.

For insulin, it was beneficial that the samples had been examined by a clinically approved ELISA kit prior to being analyzed by the LC-MS method [30], as a suitable calibration concentration range could easily be selected. For the other hormones, somatostatin-14 and glucagon, a gold standard for determination of the concentrations have not yet been established [31–33]. A calibration concentration range was selected without prior information about expected concentration of analytes in the sample, the concentration in the samples were found to be below or around the lowest concentration calibration standard (0.25 pg/µL). Therefore, the samples collected with 20 mM glucose and 30 mM KCl were reexamined with a calibration from 0.1 pg/µL to 3 pg/ µL for stomatostatin-14, and 0.05 pg/µL to 3 pg/µL for glucagon, including QCs at 0.4 pg/µL. The concentration for somatostatin-14 could now be determined and was found to be 0.27 pg/µL (RSD = 20%, n = 4, N = 1) in the samples collected with KCl. For glucagon, the concentration was determined with the new calibration curve to be 0.31 pg/µL (RSD = 18%, n = 4, N = 1), which was not significantly different form the concentration determined with the first calibration curve based on an independent two sample t-test (95 % confidence). The QCs examined at 0.4 pg/µL was all within 10% relative error for somatostatin-14 (N = 3), and 11% relative error for glucagon (N = 3).

The carry-over was less than 1% for all of the analytes and the internal standard, shown in a representative chromatogram from a blank injection of 0.5% FA in Krebs buffer following injections of the standards used to establish the curve in **Figure 4E**. In addition, the carry-over was equal to less than 20% of the peak area obtained in the calibration standard with the smallest concentration of the analytes, giving a lower limit of quantification of 0.2 pg/µL for human insulin, 0.1 pg/µL for somatostatin-14, and 0.05 pg/µL for glucagon. The retention time variation was less than 0.2% RSD (N = 26) for all of the analytes and the internal standard.

A challenge when examining supernatant collected from SC-islets after exposure to different levels of glucose and KCl, is the change in the sample matrix. In this study, Krebs buffer without glucose or KCl was used as the solution for preparation of the calibration standards. Possible effects of glucose and KCl in the samples has not been examined. The ratio of the quantifier and qualifier transitions obtained for each hormone is shown in **Figure 4F**. For bovine and human insulin, the quantifier/qualifier ratio was not significantly different in samples with different amounts of glucose or KCl (determined with one-way ANOVA). In addition, for all analytes and bovine insulin, an independent two sample t-test, at 95% confidence, confirmed there was no significant difference in the quantifier/qualifier ratio obtained in the samples compared with the quantifier/qualifier ratio obtained in the calibration standards and QCs. The identification and quantification in the RPLC-MS/MS method is secured by negligible variance in retention time and quantifier/qualifier ratios, negligible levels of carry-over (< 1%), and sufficient precision and accuracy.

In conclusion, the RPLC-MS/MS method demonstrates sufficient detection limits for determination of insulin secretion in the supernatant of SC-islets (n = 30). In addition, we show that the SC-islets obtained through our differentiation protocol [18], have obtained a glucose dependent insulin secretion in response to 2 mM and 20 mM glucose. The method can also determine the production of somatostatin-14 and glucagon in 20 mM glucose and 30 mM KCl. However, better sensitivity is needed to determine secretion of somatostatin-14 and glucagon in SC-islets (n = 30) stimulated by glucose alone.

### 3.4 Insulin could be determined in background secretion from a small number of stem cell-derived islet organoids cultivated in a pump-less, recirculating organ-on-a-chip device

Hormone secretion from isolated mouse or human islets-on-chip has been determined with the use of e.g. luminescent immunoassay (AlphaLISA) [34], or ELISA [35]. However, to the authors’ knowledge, the combination of human SC-islets, organ-on-a-chip device, and hormone secretion determination with LC-MS has not been previously reported. As a proof-of-concept for combining RPLC-MS/MS determination of intact hormones from SC-islets cultured on an organ-on-chip device, we wanted to examine background secretion of the hormones. In addition, we wanted to show that the RPLC-MS/MS method was versatile concerning the applied sample matrix and therefore did not change the islet maturation cell medium (will be referred to as cell medium) with Krebs buffer for this experiment.

The background secretion over 24 h (in cell medium containing 5.5 mM glucose), from 3-6 SC-islets cultivated on-chip in a rOoC device was compared to 14-19 SC-islets cultivated in a standard cell culture well-plate.

To avoid introducing variation in the quantification, bovine insulin was not applied as an internal standard during the determination of human insulin in cell medium, due to a higher variation in the peak area (RSD > 7 %, N = 6) in cell medium compared to Krebs buffer (RSD < 5%, see **Section 3.2**). At day 5 on the rOoC, an average of 5 pg/µL of insulin was secreted per SC-islet (RSD = 52%, n = 4, N = 1), while on day 7 an average of 3 pg/µL of insulin was secreted per SC-islet (RSD = 30%, n = 4, N = 1). Similarly, on the well-plate at day 5, an average of 7 pg/µL of insulin was secreted per SC-islet (RSD = 24%, n = 4, N = 1), while on day 7 an average of 3.9 pg/µL of insulin was secreted per SC-islet (RSD = 7%, n = 4, N = 1). An independent two sample t-test, at 95% confidence, confirmed there was no significant difference in insulin secretion per SC-islet on the rOoC compared to the cell culture well-plate. The high variation in the determination of insulin secretion per SC-islet on the rOoC device (54% on day 5 and 30% on day 7) compared to the 24 well-plate (24% on day 5 and 7% on day 7), can be explained by the number of SC-islets included on the different devices. There was 3-6 SC-islets on the rOoC, while there was 14-19 SC-islets included on the well-plate. Individual differences in the SC-islets may affect the reliability of the measurements when examining a small batch of SC-islets, and that the lower RSD values on the 24 well-plate indicates that a representative batch of SC-islets should be closer to 20 individual SC-islets.

Concerning background secretion of somatostatin-14 and glucagon in 3-6 SC-islets cultivated on rOoC device, the detection limits was not sufficient for quantification of secretion from limited number of SC-islets stimulated by only glucose, for details see **SI-3.** We were able to determine secretion of glucagon in the samples collected from 14-19 SC-islets on the cell culture well-plate.

The composition of the SC-islets was confirmed with flow cytometry quantification and immunostaining (**Figure 5A**), for details see **SI-4**. The SC-islets consisted of >66% insulin-positive cells, >17% somatostatin-positive cells, and >22% glucagon-positive cells, where >95% of the cells were endocrine cells (i.e. cells which can secrete hormones). The multicellular SC-islets have a composition, which is similar to the distribution of the cell types in human islets [13, 15].

**Figure 5:**
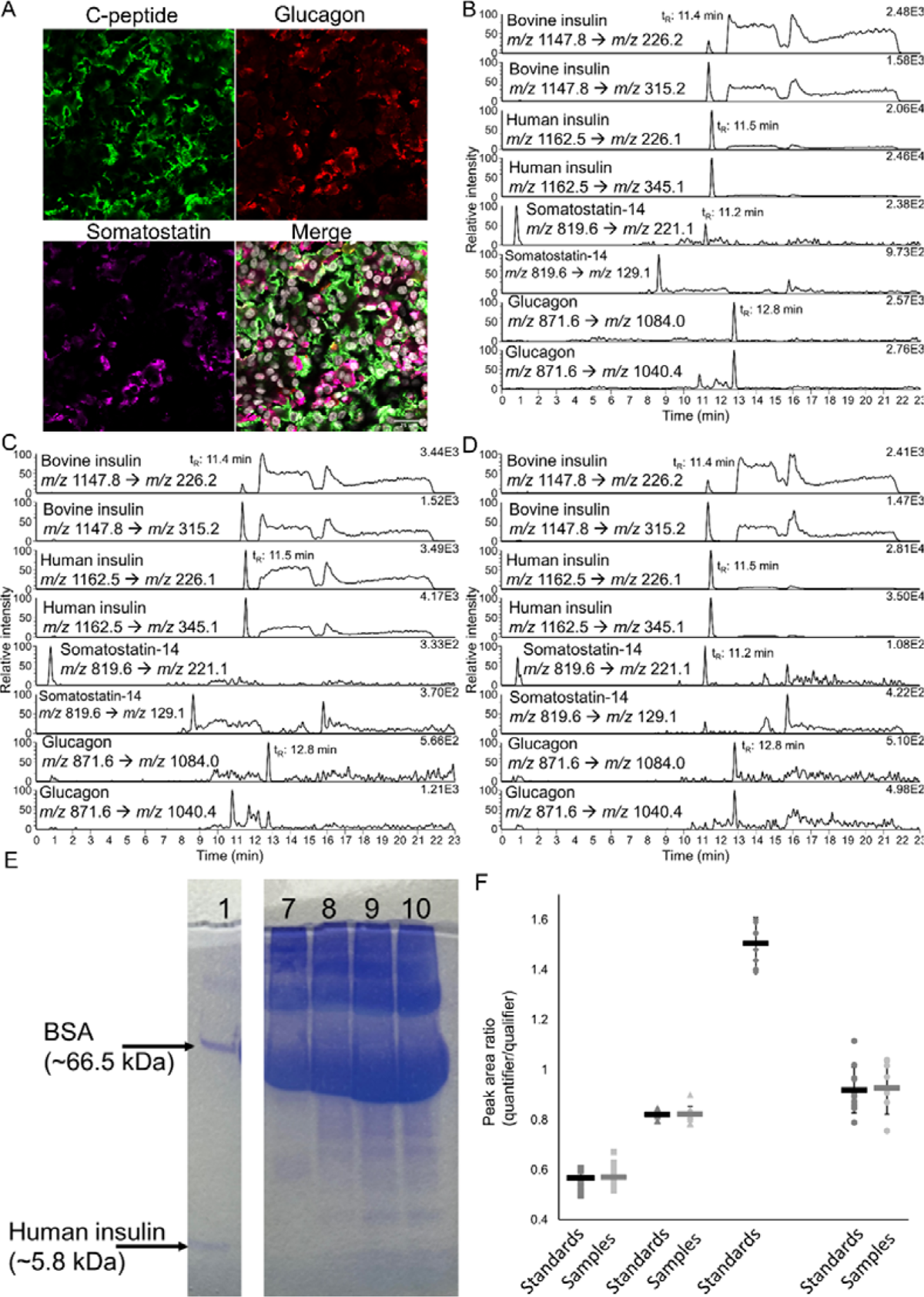
(**A**) Representative immunostaining images of SC-islets stained for C-peptide (green), glucagon (red), somatostatin-14 (magenta), and for cell nuclei with Hoechst 333242 (white). Scale bar = 25 µm. Representative chromatograms obtained in SRM mode of: Supernatant from SC-islets cultivated (**B**) in a 24 well-plate and (**C**) on the rOoC device, and (**D**) calibration standard with 100 pg/µL human insulin, 0.1 pg/µL somatostatin-14, and 0.1 pg/µL glucagon with 5 pg/µL bovine insulin in 1.0% FA in cell medium. The first transition for each hormone is the quantifier, while the second transtion is the qualifier. (**E**) Coomassie blue stained protein bands found by gel electrophoresis in the following samples: (**1**) 10 ng/µL og human insulin and 10 ng/µL of BSA in water, (**7**) 1.0% FA in cell medium, (**8**) pooled supernatant collected on day 5 from SC-islets cultured in a 24 well-plate, (**9-10**) two replicates of supernatant from SC-islets incubated on rOoC collected on day 5. The picture of the gel has been cropped, where Lane 2-6 is not included, however, the picture of the gel is provided in raw format in **SI-5**. (**F**) The ratio of quantifier/qualifier transitions for bovine insulin (squares), human insulin (triangles), somatostatin-14 (diamonds), and glucagon (circle) obtained for calibration standards and QCs (dark grey), and samples (light grey). Both transitions of somatostatin-14 were not detected in any of the examined sample.

To summarize, determination of secreted insulin from limited number SC-islets (n < 6) cultivated in cell medium on a rOoC device was possible with the applied RPLC-MS/MS method. It was found that the insulin secretion in the SC-islets was not significantly different on the rOoC device compared to the secretion occurring on a standard cell culture well-plate.

#### 3.4.1 Discussion: concerning challenges with cell medium, the organ-on-chip device, and the applied liquid chromatography mass spectrometry method

In the present method an interfering peak (eluting after 13 min in standard solutions in cell medium) is eluting closer to the analytes and co-elute with glucagon in cell medium incubated with SC-islets on well-plate (**Figure 5B**) and on rOoC device (**Figure 5C**). In comparison, there is a baseline separation of the analytes and the interfering peak in the standard solution prepared with fresh cell medium spiked with the analytes and internal standard (**Figure 5D**). The separation in the samples may be affected by other sample matrix components introduced following incubation of the SC-islets on the 24 well-plate or in the rOoC device. By gel electrophoresis, it was possible to confirm the presence of varius proteins in the size range between human insulin and BSA (See Lane 1, **Figure 5E**) in supernatant collected from SC-islets that were either cultivated in a 24 well-plate (Lane 8) or in the rOoC device (Lane 9 and 10). The observations suggest that the extra sample matrix components are released into the cell medium by the SC-islets or the extracellular matrix used for embedding the SC-islets on the two devices. Indeed, in cell medium neither incubated with SC-islets nor been in contact on either of the devices, significantly less protein bands were visible in the same size range. Gel electrophoresis was also used to compare cell medium with Krebs buffer concerning protein content, showing that there were more sample matrix components present in cell medium compared to Krebs buffer, see **SI-5** for more details.

It is worth nothing that during preliminary examination of the rOoC device with Krebs buffer, there was a significant loss of human insulin following incubation on the rOoC device compared to standard cell culture well-plates, see details in **SI-6**.

In the current experiment, the reliability of the determination of hormones with the RPLC-MS/MS method was not affected by the changes to the separation due to the presence of additional sample matrix components. The identification was secured by a stable ratio of the quantifier and qualifier transitions obtained for each hormone in calibration standards and sample is shown in **Figure 5F**, as there was no significant difference in the ratio obtained in calibration standards and samples. In addition, the retention time of each hormone varied ≤ 0.5% RSD (N = 30) and the carry-over was < 2% for glucagon, < 0.5% for human insulin, and < 0.1% carry-over for bovine insulin and somatostatin-14. Additionally, all QCs (examined before, within and after the sample-set) at 25 pg/µL of human insulin, and 0.8 pg/µL of somatostatin-14 and glucagon were determined within ± 15 relative error (N = 5). The only exception was for the determination of glucagon, where the relative error was 17% in the second replicate of the QC.

To summarize, the RPLC-MS/MS offers reliable determination of insulin, somatostatin-14 and glucagon in a complex matrix (cell culture supernatant in this study) without use of internal standard, with negligible variation in retention time, repeatable quantifier/qualifier transition ratios, and negligible levels of carry-over. With the growing complexity of the cell medium samples following incubation with the SC-islets embedded with extracellular matrix on the 24 well-plate and in the rOoC device, we are approaching a limit where the RPLC-MS/MS method alone is not sufficient for reliable determination. In the case of even more complex samples, the inclusion of sample preparation steps might become necessary.

## 4 Concluding remarks

The study was dedicated to combining the determination of multiple peptide hormones with liquid chromatography-mass spectrometry and SC-islets-on-a-chip. It has been shown that liquid chromatography is needed to separate the target peptides from interferences in the sample matrices. However, when the complexity of the samples grows and large amounts of proteins are present, chromatography and mass spectrometry may not be enough for successful peptide determination (e.g. urocortin-3 co-elutes with BSA), pointing to the need for the inclusion of sample preparation steps, e.g. electromembrane extraction of target peptides.

## Acknowledgements

Financial support was obtained from the Research Council of Norway through its Centres of Excellence funding scheme, project number 262613 and partly from the UiO:Life Science convergence environment funding scheme. S.R.W. is a member of the National Network of Advanced Proteomics Infrastructure (NAPI), which is funded by the Research Council of Norway INFRASTRUKTUR-program (project number: 295910).

## Conflict of interest

The authors declare no conflict of interest.

## Data Availability Statement

The data that support the findings of this study are available from the corresponding author upon reasonable request.

## Abbreviations

A_s_: Asymmetry factor

FA: Formic acid

MEM NEAA: Minimum essential medium non-essential amino acids

N: Technical replicate

n: Biological replicate

RF: Radio frequency

rOoC: Recirculating organ-on-chip

## Notes

### Competing Interest Statement

The authors have declared no competing interest.

