## Supporting Information for "Simultaneous LC-MS determination of glucose regulatory peptides secreted by stem cell-derived islet organoids"

*Supplementary materials for*: Simultaneous LC-MS determination of glucose regulatory peptides secreted by stem cell-derived islet organoids

Christine Olsen^1,2^, Chencheng Wang^2,3^, Aleksandra Aizenshtadt^2^, Shadab Abadpour^2,3^, Elsa Lundanes^1^, Frøydis Sved Skottvoll^4^, Alexey Golovin^5^, Mathias Busek^2^, Stefan Krauss^2^, Hanne Scholz^2,3^, and Steven Ray Wilson^2,3*^

^1^ Department of Chemistry, University of Oslo, Blindern, Oslo, Norway

^2^ Hybrid Technology Hub-Centre of Excellence, Institute of Basic Medical Sciences, Faculty of Medicine, University of Oslo, Oslo, Norway

^3^ Department of Transplant Medicine and Institute for Surgical Research, Oslo University Hospital, Oslo, Norway

^4^ Department of Smart Sensors and Microsystems, SINTEF Digital, Oslo, Norway

^5^ Department of Immunology and Transfusion Medicine, Oslo University Hospital, Oslo, Norway

*Correspondence should be addressed to the following author(s):

Prof. Steven Ray Wilson

Department of Chemistry

University of Oslo

Postboks 1033, Blindern 0315 Oslo

.

### 1 Significantly increased sensitivity at higher collision gas pressures for insulin and glucagon

The LC-MS method applied in this study, had previously been optimized for human insulin where a collision gas pressure of 2.5 mTorr had been selected based on reduced variance of fragmentation based on direct injection [1]. With the introduction of other peptides, the collision gas pressure was reexamined to see if the obtained peak areas and sensitivity of the method could be increased. The average peak area of human insulin obtained at 4.0 mTorr was 11663 with a RSD of 12% (n = 3), while at 2.5 mTorr the average peak area was 8746 with a RSD of 8% (n = 3). For glucagon the average peak area was 88028 (8 % RSD, n = 3) and 64654 (3% RSD, n = 3) for 4.0 mTorr and 2.5 mTorr, respectively. An independent two sample t-test, at 95% confidence, showed that the peak areas obtained for human insulin and glucagon were significantly higher for 4.0 mTorr collision gas pressure compared to 2.5 mTorr, while the variances at each gas pressure were not significantly different. For somatostatin-14 there was no significant difference depending on applied collision gas pressure; the peak area at 4.0 mTorr was 35886 (12% RSD, n = 3), while the peak area was 33708 (5% RSD, n = 3) at 2.5 mTorr. Based on the increased sensitivity for human insulin and glucagon, a collision gas pressure of 4.0 mTorr was applied in the following experiments.

### 2 Estimation of gradient delay

During gradient optimization, the gradient delay was examined by running the analysis of the peptides in non-retained conditions, as the elution time of the peptides would help in determining at which %B the target analytes eluted in the original shallow gradient from 1% B to 60% B in 8 min (an increase of 7.4% B/min). The system delay was estimated by analyzing a 20 pg/µL peptide mix (consisting of bovine insulin, human insulin, somatostatin-14, and glucagon) in 0.5% FA in Krebs buffer. A gradient starting at 60% B running up to 90% B was applied, and the obtained peptide peaks co-eluted at approx. 6.5 min (N = 2), see **Figure S1**. With an elution time of 11.3 min for bovine insulin in water, the expected %B at the elution was calculated by: (11.3 min – 6.5 min)*7.4% B = 35.5% B. The gradient was step-wise optimized based on this estimation, to separate the target analytes from the interferences in Krebs buffer and cell medium.

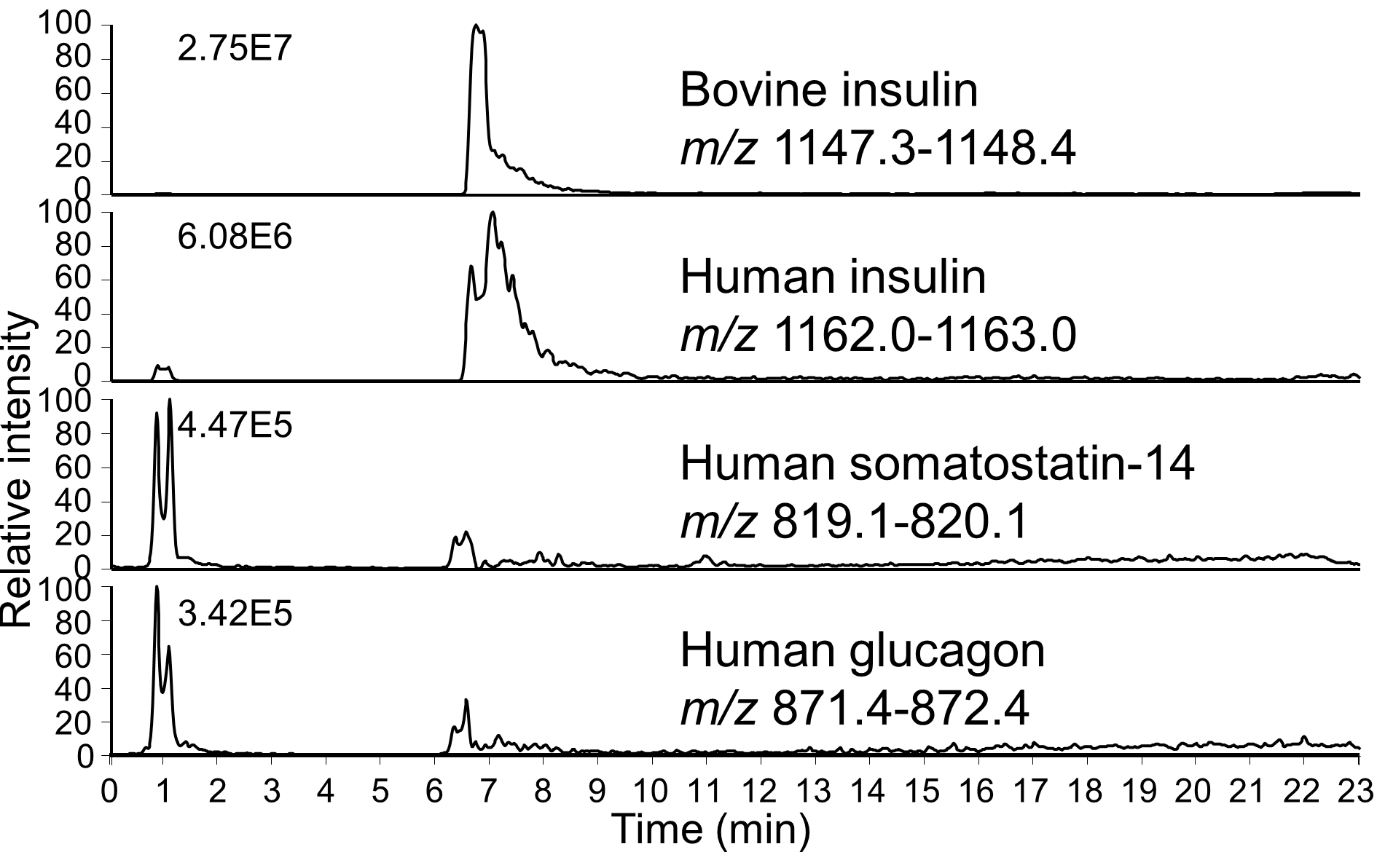

**Figure S1**: Representative chromatogram obtained in fullscan mode for 20 pg/µL of human insulin, somatostatin-14, glucagon, and 5 pg/µL of bovine insulin in 0.5% FA in Krebs buffer. The peaks co-eluted at 6.5 min when running the chromatography at non-retained conditions by starting the gradient at 60% B.

### 3 Comments concerning attempted quantification of background secretion of somatostatin-14 and glucagon from SC-islets during cultivation

During cultivation on a well-plate or a rOoC device, the SC-islet organoids are maintained with cell medium containing 5.5 mM glucose. The glucose should stimulate the SC-islets to secrete peptide hormones, including insulin (discussed in main document in Section 3.4), somatostatin-14 and glucagon.

For somatostatin-14, the peak area obtained in supernatant from the well-plate at day 5 was 647 (RSD = 40, n = 3, N = 1), and 786 (RSD = 29, n = 3, N = 1) on day 7 (representative chromatogram in **Figure S2A**), while on the rOoC at day 5 the peak area was 522 (RSD = 30, n = 3, N = 1), and 409 (RSD = 8, n = 3, N = 1) on day 7 (representative chromatogram in **Figure S2B**), which is in the range of the estimated limit of detection. By comparing the peak areas in the samples with the peak area of 701 obtained from 0.1 pg/µL of somatostatin-14 (**Figure S2C**), it is highly probable that the SC-islets do secrete somatostatin-14. However, as seen in the chromatograms, the qualifier transition for somatostatin-14 (*m/z* 819.6 to *m/z* 129.1) was not detected in the samples from the well-plate or the rOoC, meaning the identification could not be confirmed.

Only glucagon was quantified in the secretion from the SC-islet organoids on the well-plate with an average of 0.13 pg/µL of glucagon per SC-islet on day 5 (RSD = 15%, n = 4, N =1) and an average of 0.025 pg/µL of glucagon per SC-islet on day 7 (RSD = 27%, n = 4, N = 1). On the rOoC, the average peak area for glucagon was 4907 at day 5 (RSD = 25%, n = 4, N =1), and 3492 on day 7 (RSD = 21%, n = 4, N =1), which was in the same area as the estimated limit of detection of 0.1 pg/µL glucagon in cell medium, which gave a peak area of 4016, shown Figure 5C.

In conclusion, a combination of better sensitivity and a larger batch of SC-islet organoids is needed to be able to determine glucose stimulated secretion of somatostatin-14 and glucagon from SC-islet organoids.

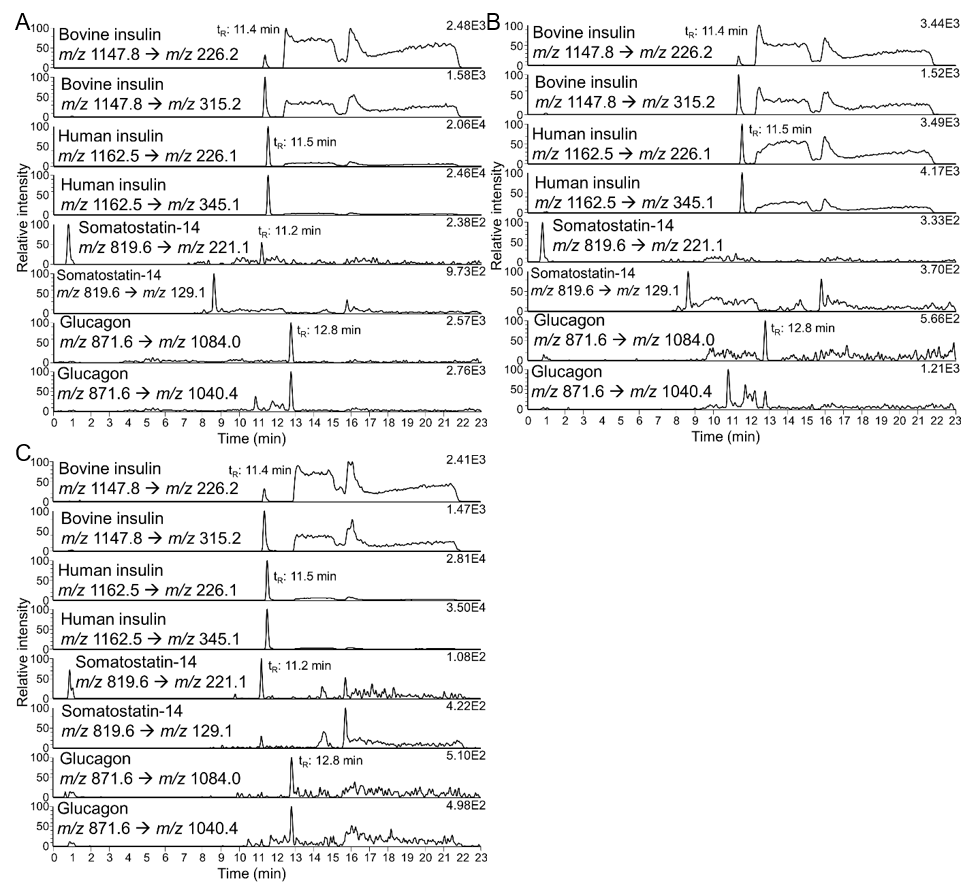

**Figure S2**: Representative chromatograms obtained in SRM mode of: Supernatant from SC-islet organoids cultivated (**A**) in well-plate and (**B**) on rOoC device, and (**C**) calibration standard with 100 pg/µL human insulin, 0.1 pg/µL somatostatin-14, and 0.1 pg/µL glucagon with 5 pg/µL bovine insulin in 1.0% FA in cell medium. The first transition for each hormone is the quantifier, while the second transtion is the qualifier.

### 4 Characterization of the cell composition in stem cell-derived islets

The SC-islets were further characterized with flow cytometry and immunofluorescence staining to confirm the presence of somatostatin and glucagon-producing cells. Primary antibodies including human C-peptide (Developmental studies Hybridoma Bank, University of Iowa, IA, USA), glucagon (GCG, Sigma Aldrich, MO, USA), somatostatin (SST, Santa Cruz, CA, USA) and chromogranin A (CHGA, Novus Biologicals, Centennial, CO, USA) represent insulin-producing cells, glucagon-producing cells, somatostatin-producing cells, and pan-endocrine cells in the organoids respectively [2]. Representative flow cytometry quantification shows that > >95% of cells are endocrine cells (Q2 and Q3, **Figure S3A-II**), and >66% of cells are insulin-producing cells (Q1 and Q2, **Figure S3A-II**). While there are >22% of cells are glucagon-producing cells (Q9 and Q10, **Figure S3A-IV**) and >17% somatostatin-producing cells (Q5 and Q6, **Figure S3A-VI**). None of the mentioned cell types were found in negative controls, **Figure S3A-I, III, and V**). SC-islets were fixed with 4% PFA at room temperature for 30 min and embedded for cryosection. Section slides immunofluorescence staining corresponded with flow cytometry results, showing that the majority of the cells are human C-peptide-positive cells, and there is a large presence of somatostatin-positive cells and glucagon-positive cells (**Figure S3B**).

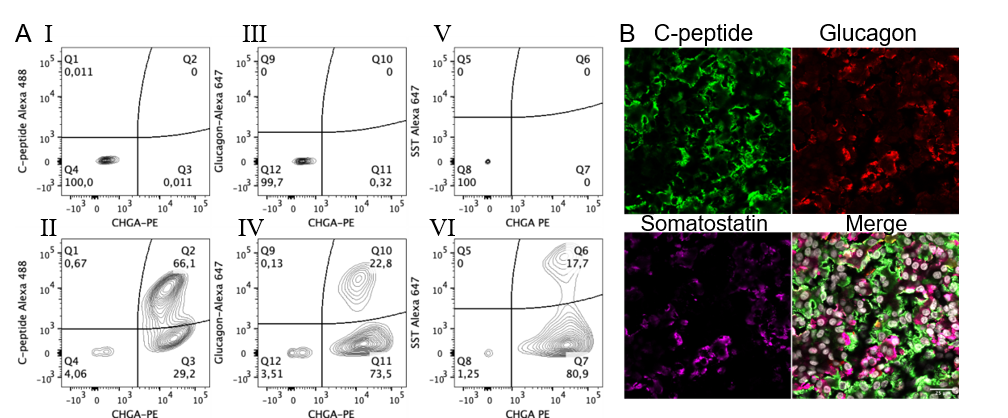

**Figure S3**: Representative flow cytometry quantification (%) of dispersed SC-islets stained for (I-II) C-peptide and chromogranin A (CHGA), (III-IV) glucagon and CHGA, and (V-VI) somatostatin-14 and CHGA. (I, III, and V) Negative control for flow cytometry quantification. (**C**) Representative immunostaining images of SC-islets stained for C-peptide (green), glucagon (red), somatostatin-14 (magenta), and merge including staining for cell nucleus with Hoechst 333242 (white). Scale bar = 25 µm.

### 5 Gel electrophoresis

Krebs buffer and cell medium have proved to be difficult matrices to work with for obtaining sufficient sensitivity in the determination of small peptide hormones and separation of the target peptide hormones from interferences found in the matrices. By comparing the obtained protein bands found in Krebs buffer (Lane 2, **Figure S4**), and cell medium (Lane 7), to a water standard containing a mixture of 10 ng/µL of human insulin and 10 ng/µL of bovine serum albumin (Lane 1), there is clearly more proteins present in Krebs buffer and cell medium.

We also examined if there were any changes to Krebs buffer following incubation on a well-plate (Lane 3) and incubation on a rOoC device (two replicates in Lane 4 and Lane 5), however, no changes were detectable with gel electrophoresis and Coomassie blue staining of proteins, see **Figure S4**.

Matrix matching in calibration standards to be applied for determination with LC-MS is of great importance. In this study, calibration standards were prepared from Krebs buffer and cell medium, however, when secreted peptides from stem cell-derived islet (SC-islet) organoids is of interest, we are unable to match the exact composition of the matrix following incubation of SC-islets. In the case of Krebs buffer, the initial composition of Krebs buffer (Lane 2, **Figure S4**) cannot be differentiated by gel electrophoresis from the composition of the supernatant collected after incubation of SC-islets in Krebs buffer with 20 mM glucose and 30 mM KCl (Lane 6). However, for cell medium, there are several other proteins bands visible in the range between the band originating from human insulin and bovine serum album (Lane 1) in supernatant collected from SC-islets incubated on well-plate (Lane 8) and on rOoC device (two replicates Lane 9 and 10), compared to cell medium (Lane 7) which has not been incubated with SC-islets.

The protein amount, and subsequently the need for separation of target hormone peptides from potential interferences is much larger in Krebs buffer compared to water, and even more important for cell medium, and in supernatant collected after incubation of SC-islet organoids.

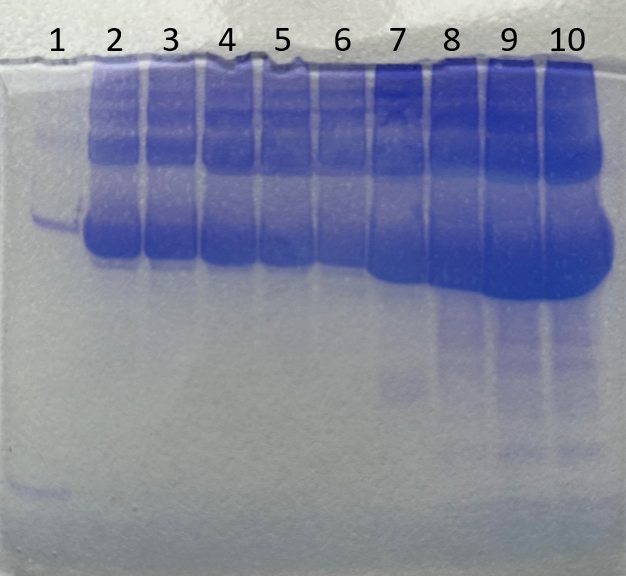

**Figure S4**: Stained protein band after by gel electrophoresis of the following samples: (**1**) 10 ng/µL og human insulin and 10 ng/µL of bovine serum albumin in water and (**2**) 0.5% formic acid in Krebs buffer. Krebs buffer incubated on: (**3**) well-plate and (**4-5**) two replicates of krebs buffer incubated on rOoC device. (**6**) Supernatant collected from SC-islet organoids incubated in Krebs buffer with 20 mM glucose and 30 mM KCl. (**7**) 1.0% formic acid in cell medium, (**8**) pooled supernatant from SC-islet organoids incubated on well-plate collected on day 5, (**9-10**) two replicates of supernatant from SC-islet organoids incubated on rOoC collected on day 5.

### 6 Significant loss of insulin in Krebs buffer incubated on a recirculating organ-on-a-chip device compared to static incubation on cell culture well-plates

Prior to analysis of SC-islet organoids on a microfluidic chip, we examined whether the measurements would be affected by non-defined adsorption of the hormones on the applied device. The rOoC device has been shown suitable for long-term cultures of human stem cell-derived liver organoids and is made out of PMMA [3]. Based on previous experience, human insulin displays a varied degree of non-defined adsorption to a large variety of tubing applied on an LC-instrumentation [1]. Therefore, we found it useful to examine if the PMMA surface in the chip would affect the recovery of our target hormones, and human insulin was used as a model analyte.

After 20 h incubation of 2 ng/µL of human insulin in Krebs buffer on the PMMA rOoC, an average of 1.58 ng/µL human insulin (79% recovery, RSD = 4%, n_r_ = 3 rOoC loops, N = 1) was recovered. In comparison, an average of 2.09 ng/µL human insulin (104% recovery, RSD = 3%, n_w_ = 3 wells, N = 1) was recovered after static incubation on a well-plate and an average of 2.12 ng/µL human insulin (106% recovery, RSD = 3%, n_a_ = 3 aliquots, N = 1) was recovered in aliquots stored in the freezer. It should be noted that in a fourth replicate on the rOoC the recovered amount of human insulin was 2.3 ng/µL, however, the replicate was rejected based on Grubbs’ test for outliers.

Based on one-way ANOVA, at least one of the recovered concentrations of human insulin was significantly different from the others. It was determined by Fisher’s least significant difference test, that there was no significant difference between the recovered concentration of human insulin on the well-plates and the control stored in the freezer. However, the recovered human insulin concentration from the rOoC was significantly different from both the concentrations recovered on the well-plate and in the control.

With a significant loss of 21% of human insulin after incubation on the rOoC in Krebs buffer, the PMMA surface on the rOoC might have to be modified to reduce adsorption of human insulin [4].
